# Impact of Peptide Initial Configuration and Membrane Composition on Melittin’s Pore-Forming Ability under Unbiased All-Atom Molecular Dynamics Simulations

**DOI:** 10.1101/2025.05.28.656502

**Authors:** Zhihao Zhao, Yilin Guo, Jiaxiang Cai, Peilin Xie, Jiahui Guan, Lantian Yao, Yulan Liu, Chia-Ru Chung, Tzong-Yi Lee, Ying-Chih Chiang

**Author notes:** Contributing authors. These authors contributed equally to this work (Zhihao Zhao, Yilin Guo, Jiaxiang Cai).

## Abstract

The rising challenge of antimicrobial resistance has accelerated the search for alternative therapeutics. Antimicrobial peptides (AMPs), a class of naturally occurring defense molecules found across diverse species, are promising candidates. Despite their potent membrane-disrupting activity, the atomic details of the pore formation process remain insufficiently understood. In this study, we employed all-atom molecular dynamics (MD) simulations to investigate the pore formation process of melittin under different initial configurations. Simulations were conducted using three different membrane systems: a pure POPC bilayer, a mammalian membrane model (DOPC:Cholesterol, 9:1), and a bacterial membrane model (DOPE:DOPG, 3:1). For each system, we examined four different starting configurations, in which six melittin peptides were arranged in a star-like pattern. Our results demonstrated that the pore formation process strongly depends on the initial peptide configuration. In one specific initial arrangement (termed as Conf.I), pore formation consistently occurred within 100 nanoseconds, regardless of membrane composition. Furthermore, the simulations revealed that pore formation was more challenging in the mammalian membrane model and even more so in the bacterial membrane model, in comparison with the pure POPC bilayer. These findings are in line with previously reported minimum inhibitory concentration (MIC) and the 50% hemolysis concentration (HC_50_) of melittin in the literature. Additionally, we identified lysine-7 (K7) as the key residue in determining whether a stable pore can form. In configurations where the K7 side chain formed electrostatic interactions with the phosphate group of a lipid, melittin were anchored to the membrane surface, thereby preventing pore formation. In contrast, simulations of melittin mutants K7A and K7Q showed no such anchoring effect, and thus pore formation was possible in multiple initial configurations. Notably, the K7Q mutation showed a preference for pore formation in bacterial membranes over mammalian membranes, suggesting that reducing toxicity while maintaining antimicrobial efficacy is possible.

## 1 Introduction

Since Alexander Fleming’s discovery of penicillin in 1928, antibiotics have served as the primary defense against bacterial infections [1]. However, widespread misuse and overprescription have accelerated the emergence of antibiotic resistance [2]. The World Health Organization estimates that antimicrobial resistance could contribute to approximately 10 million deaths annually by 2050 [3, 4], underscoring the urgent need for alternative therapeutic strategies. Antimicrobial peptides (AMPs) have emerged as promising candidates to address this challenge [5–7]. These peptides exhibit broad-spectrum antimicrobial activity and possess diverse functionalities, including antiviral, antifungal, anticancer, and immunomodulatory properties, making them viable alternatives to conventional antibiotics [8–11]. Currently, the most comprehensive AMP database, dbAMP 3.0, catalogs 35,518 AMPs and 2,453 antimicrobial proteins sourced from 3,534 species, with ongoing updates and expansion [12]. The rapid expansion of the AMP database highlights the significance of AMP development. Recent advancements in artificial intelligence (AI) have fundamentally transformed the research and development of AMPs [13, 14]. Computational frameworks, such as Multi-Conditional Generative Adversarial Networks [13] and Transformer-based neural network architectures [15], now achieve prediction accuracies exceeding 90%. While sequence-based prediction algorithms effectively identify statistically significant residues, they lack the ability to provide atomic-level insights into the mechanisms of action of AMPs. This is particularly crucial when addressing questions such as why one AMP sequence exhibits higher activity against bacteria than another, or whether a specific sequence demonstrates greater toxicity than alternatives.

Conventional experimental techniques so far are inadequate for capturing the transient pore formation process at atomic resolution. As a result, insights into the mechanisms of the pore formation process and the underlying peptide-lipid interactions, largely rely on molecular dynamics (MD) simulations [16, 17]. Recent advances in computational power have enabled millisecond-long MD simulations of systems containing billions of atoms, which allows researchers to employ MD techniques to simulate critical AMP mechanisms such as membrane pore formation, lipid disruption, and transmembrane insertion [18, 19]. However, in conventional (unbiased) all-atom MD simulations, peptides typically remain on the membrane surface, i.e. so called “surface-bound state” (S-state), due to the high energy barriers associated with peptide insertion. Consequently, spontaneous pore formation was often not observed in standard MD simulations [20–23]. To simulate pore formation process caused by AMPs, various approaches have been employed. For example, Ulmschneider et al. demonstrated that WALP peptides can unfold and insert into DPPC/DMPC lipid bilyaers at 353 K [24]. Subsequently, simulations of maculatin at even higher temperatures (363.15-423.15 K) revealed the spontaneous formation of membrane channels [25]. These channels were transient, functional pores formed by various number of helices, structurally resembling transmembrane proteins. Further developments included the use of shorter lipids to enable simulations at reduced temperatures (343.15 K) [26]. Alternatively, coarse-grained modeling via the Martini force field has also been employed to investigate the pore formation process. Examples include studies on the membrane poration kinetics of melittin [27] and the localization preference of AMPs on membranes composed of different lipid phases [28]. The latter study observed the spontaneous aggregation of magainin-2 into star-like assemblies on the membrane surface, which was subsequently followed by pore formation. Notably, similar configurations (star-like assemblies) were observed also in other Martini simulations [29, 30]. Interestingly, Miyazaki et al. recently observed a rapid poration by melittin using conventional all-atom MD simulations. Their simulations were conducted at physiological temperature, and initiated from a specific initial configuration, viz six peptides were initially arranged in a star-like configuration with their N-termini oriented toward the center [31]. Under these conditions, melittins were partially inserted into the upper leaflet of the membrane. Subsequently, through collective insertion of melittins, spontaneous pore formation was observed within 100 nanoseconds. These findings suggest that poration may result from the collective motion of AMP aggregates. Following this idea, we investigate the impact of different initial configurations of melittin assemblies, as well as variations in membrane composition, on poration dynamics.

## 2 Method

### 2.1 Simulation Systems

A total of 24 systems were constructed to investigate pore formation under various conditions (Table 1). Each system contained a lipid bilayer with different membrane compositions: pure POPC, DOPC:Cholesterol at a 9:1 ratio, representing a mammalian cell membrane model, and DOPE:DOPG at a 3:1 ratio, representing a Gram-negative bacterial cell membrane model [32, 33]. All bilayers were constructed using CHARMM-GUI [34–36], consisting of 256 lipids evenly distributed between the upper and lower leaflets (128 lipids per leaflet). The systems were solvated with water molecules and salted with NaCl at a physiological concentration of 0.15 M, along with counter ions to ensure overall charge neutrality. Melittin (PDB code: 2MLT) is a 26-residue AMP with the sequence GIGAVLKVLTTGLPALISWIKRKRQQ-NH_2_. It consists of a hydrophobic N-terminal segment (residues 1-11) and a highly charged, hydrophilic C-terminal segment (residues 12-26), connected by a characteristic kink at Pro14 [37]. In this study, we also investigated two melittin mutants, K7A and K7Q, in which the lysine at position 7 was substituted with alanine and glutamine, respectively. The structures of these mutants were predicted using AlphaFold 3 [38]. As illustrated in Figure 1, the helical structure and amino acid distribution of melittin highlight its amphipathic nature, with clearly defined polar and nonpolar regions [39, 40]. Melittin carries a net positive charge of +6 and can undergo a conformational transition from a disordered random coil in aqueous environments to an amphipathic *α*-helical structure upon membrane association [41].

**Table 1.**
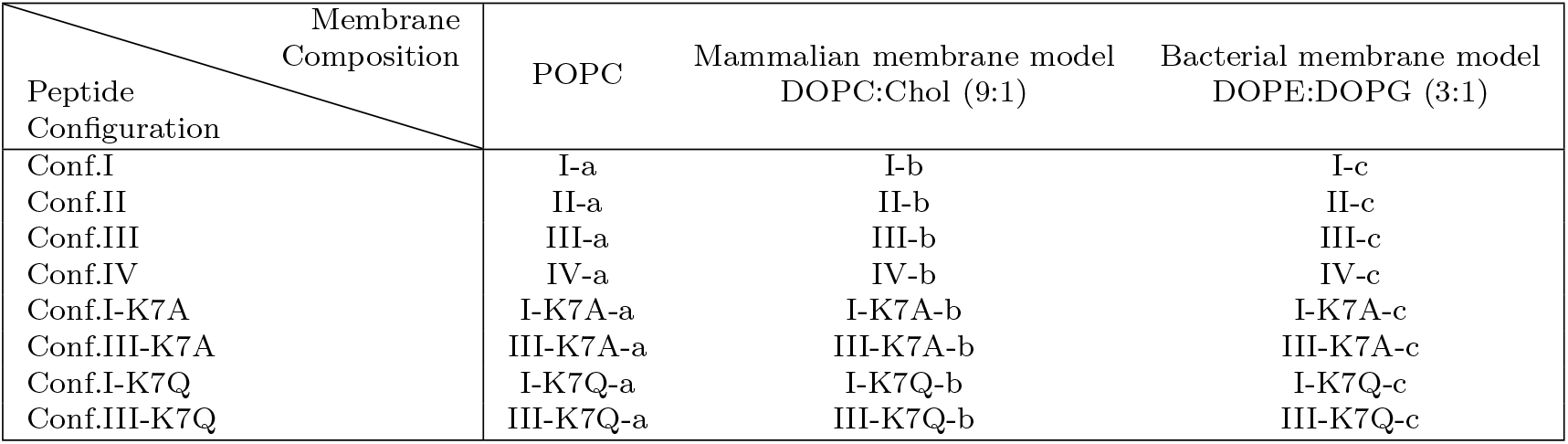
Simulated systems in this work.

| Peptide Configuration | Membrane Composition |  |  |
| --- | --- | --- | --- |
|  | POPC | Mammalian membrane model<br>DOPC:Chol (9:1) | Bacterial membrane model<br>DOPE:DOPG (3:1) |
| Conf.I | I-a | I-b | I-c |
| Conf.II | II-a | II-b | II-c |
| Conf.III | III-a | III-b | III-c |
| Conf.IV | IV-a | IV-b | IV-c |
| Conf.I-K7A | I-K7A-a | I-K7A-b | I-K7A-c |
| Conf.III-K7A | III-K7A-a | III-K7A-b | III-K7A-c |
| Conf.I-K7Q | I-K7Q-a | I-K7Q-b | I-K7Q-c |
| Conf.III-K7Q | III-K7Q-a | III-K7Q-b | III-K7Q-c |

**Fig. 1.**
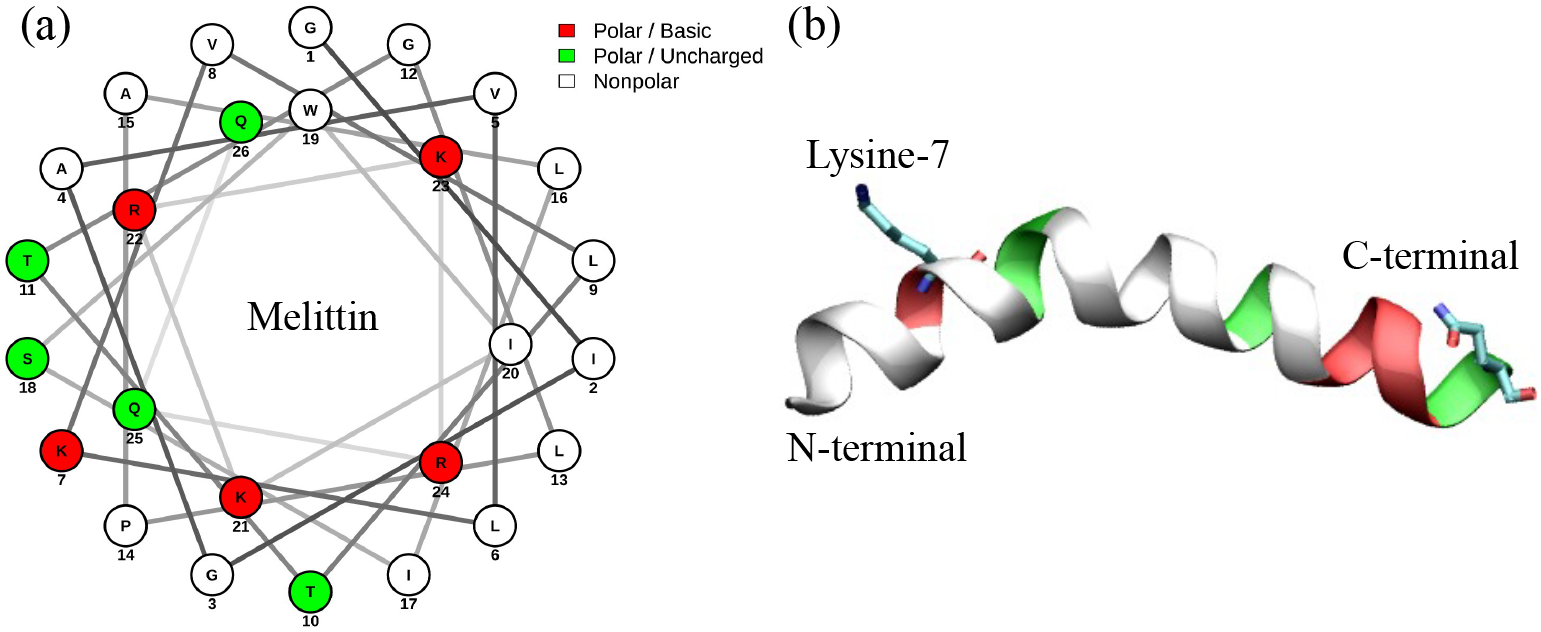
Structural representation of melittin. (a) Helical wheel projection of melittin illustrating the distribution of amino acids. The color scheme indicates residue types: red for polar/basic residues, green for polar (uncharged) residues, and white for nonpolar residues. Numbers correspond to the positions of each amino acid in the sequence. (b) Three-dimensional structure of melittin, with the helix colored according to the same scheme as in panel (a).

As shown in Figure 2, four initial configurations were:

- **Configuration I (Conf.I)**: The C-terminal segments are placed on the membrane surface, while the N-terminal segments are oriented toward the membrane center and are partially embedded in the upper leaflet. (This is the same configuration as reported in literature [31].)
- **Configuration II (Conf.II)**: Similar to Conf.I, but the peptides are shifted upward by approximately 2 Å above the membrane surface.
- **Configuration III (Conf.III)**: The C-terminal segments are pointed away from the membrane, while the N-terminal segments are positioned parallel to the membrane surface and are partially embedded in the upper leaflet.
- **Configuration IV (Conf.IV)**: Similar to Conf.III, but the peptides are shifted upward by approximately 2 Å above the membrane surface.

**Fig. 2.**
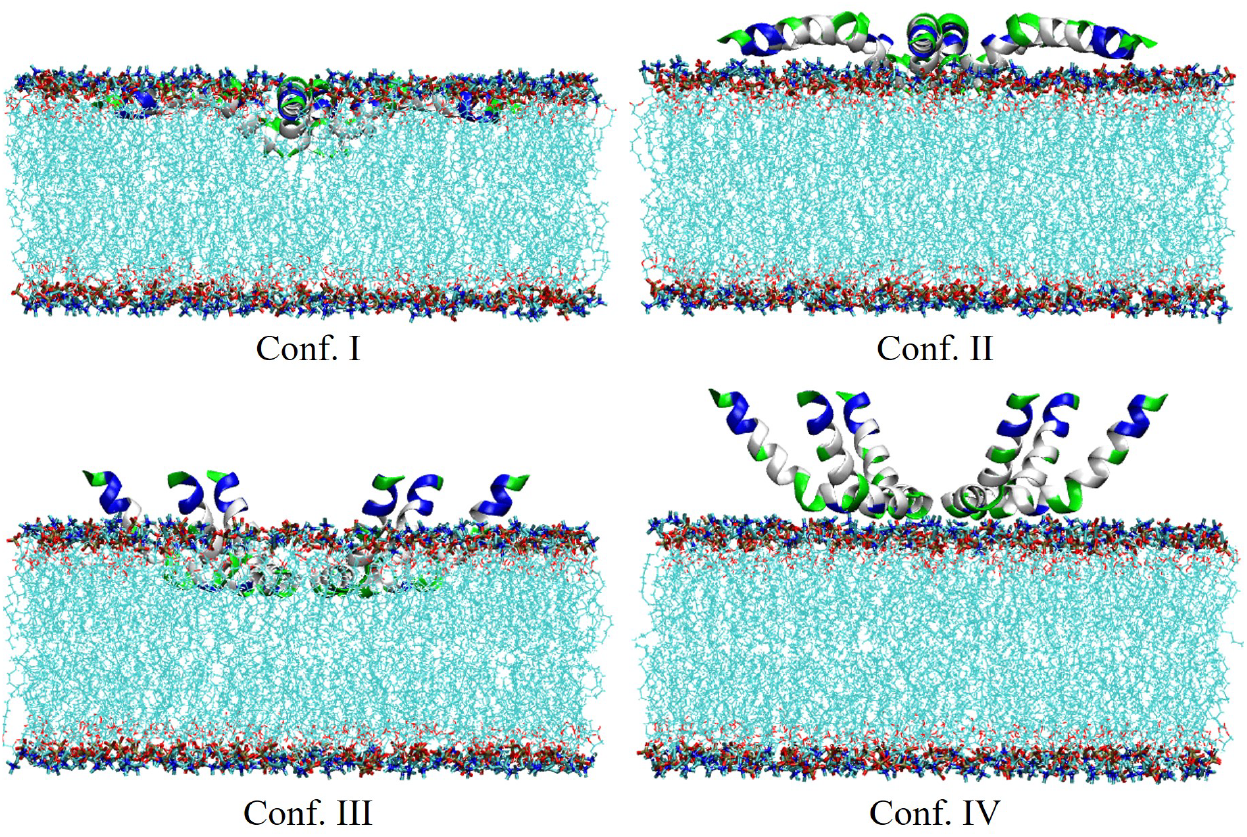
Initial configurations of melittin employed in this study. Six melittin peptides are arranged in a star-shaped assembly with their N-termini oriented toward the center. Conf.I: The C-terminal segments are placed on the membrane surface, while the N-terminal are partially embedded in the upper leaflet. Conf.II: Similar as Conf.I, but shifted upward by approximately 2 Å above the membrane surface. Conf.III: The C-terminal are pointed away from the membrane, while the N-terminal segments are placed parallel to the membrane surface and partially embedded in the upper leaflet. Conf.IV: Similar as Conf.III, but shifted upward by approximately 2 Å above the membrane surface.

Additional systems incorporating the K7A and K7Q mutations of melittin were prepared in both Conf.I and Conf.III configurations (denoted as Conf.I-K7A, Conf.III-K7A, Conf.I-K7Q, and Conf.III-K7Q) to elucidate the functional role of this specific residue. For each system, three independent simulations were performed to ensure reproducibility of the observed phenomena and statistical robustness in subsequent analyses.

### 2.2 Simulation Details

All MD simulations were performed using GROMACS 2024.2 [42, 43]. Each system was prepared through energy minimization, followed by two equilibration phases: 250 ps in the NVT ensemble and 1,625 ps in the NPT ensemble. During equilibration, restraints on protein backbone atoms, side chains, and lipid molecules were gradually lifted. Production runs were carried out using the leap-frog integrator with a 2 fs time step, each running for 750 ns, with three independent replicates per system. Pore formation events were typically observed within the first 100 ns in energetically favorable configurations, indicating that the 750 ns simulation duration was sufficient to capture pore formation propensities across different initial configurations. This timescale allows for a clear distinction between configurations that readily facilitate pore formation and those that encounter substantial energetic barriers under physiological conditions.

The CHARMM36m [44] force field was employed for peptides, lipids, and ions, while water molecules were represented using the TIP3P model. The temperature was maintained at 310 K using the velocity-rescale thermostat [45] with a coupling time constant of 1.0 ps. Pressure was controlled at 1 bar using the C-rescale barostat [46] (coupling time constant: 5.0 ps; compressibility: 4.5 *×* 10^*−*5^ bar^*−*1^) with semi-isotropic coupling to account for membrane-specific dynamics. Long-range electrostatic interactions were treated using the particle mesh Ewald (PME) method [47, 48] with a real-space cutoff of 1.2 nm. Van der Waals interactions were modeled using a force-switch modifier between 1.0 to 1.2 nm. The Verlet cut-off scheme was employed, with neighbor lists updated every 20 steps. Bond constraints involving hydrogen atoms were applied using the LINCS algorithm [49, 50]. Center of mass motion was removed every 200 fs for the solute-membrane and solvent groups independently. Trajectory coordinates were saved at 100 ps intervals for subsequent analysis.

## 3 Results and Discussion

### 3.1 Impact of Initial Configuration on Pore Formation

In POPC membrane systems, pore formation was observed exclusively in system I-a, consistent with previous findings [31, 51]. The rapid poration process began with the collective insertion of melittin peptides into the POPC bilayer, resulting in the formation of a stable transmembrane channel, as illustrated by the trajectory snapshots in Figure 3. Analysis of the peptides’ center of mass (COM) further supports this conclusion. As shown in Figure 4(a), the COM of melittin peptides along the z-axis was located between the COMs of phosphate groups in the upper and lower leaflets in system I-a, suggesting a successful pore formation. Notably, the cationic C-terminal residues (KRKR) formed a stable electrostatic interactions with the phosphate groups of POPC lipids, causing the melittins’ COM to center at about 1 nm above the membrane center (z=0). In contrast, the same analysis for the other three systems showed no peptide insertion. In particular, in panels (b) and (d), the COM of melittins was located around 5 nm above the phosphate groups of the upper leaflet, indicating that simulations initiated with peptides in the solvent (system II-a and IV-a) did not show any pore formation within 750 ns. Figure S1 provides representative frame snapshots for systems II-a and IV-a, illustrating their non-porating configurations. In system III-a, although pore formation was not observed, the melittins remained associated with the membrane surface, adopting the S-state. See Figure 4(c) the position of the peptides’ COM closely following the location of the COM of phosphorus atoms in the upper leaflet.

**Fig. 3.**
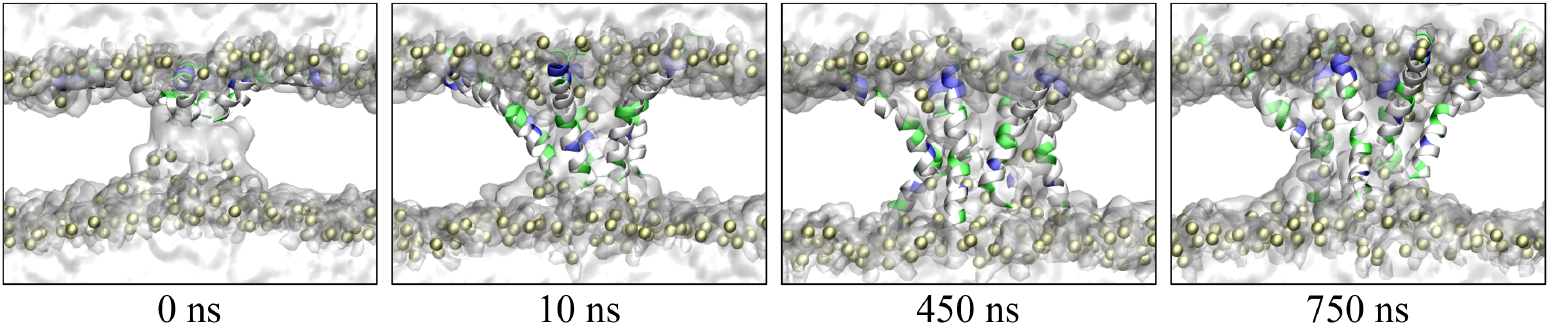
Pore formation in the POPC bilayer induced by melittin peptides in system I-a. At 0 ns, the peptides are partially embedded in the upper leaflet. Water molecules already enter the membrane center during the equilibration step. By 10 ns, membrane deformation is observed, and the peptides begin to insert into the bilayer. At 450 ns, a transmembrane channel (pore) is formed, which remains stable through the end of the simulation at 750 ns. Peptides are shown in cartoon representation, colored according to residue type. Phosphorus atoms of POPCs are displayed in van der Waals (VDW) representation, while water molecules are depicted as a transparent surface.

**Fig. 4.**
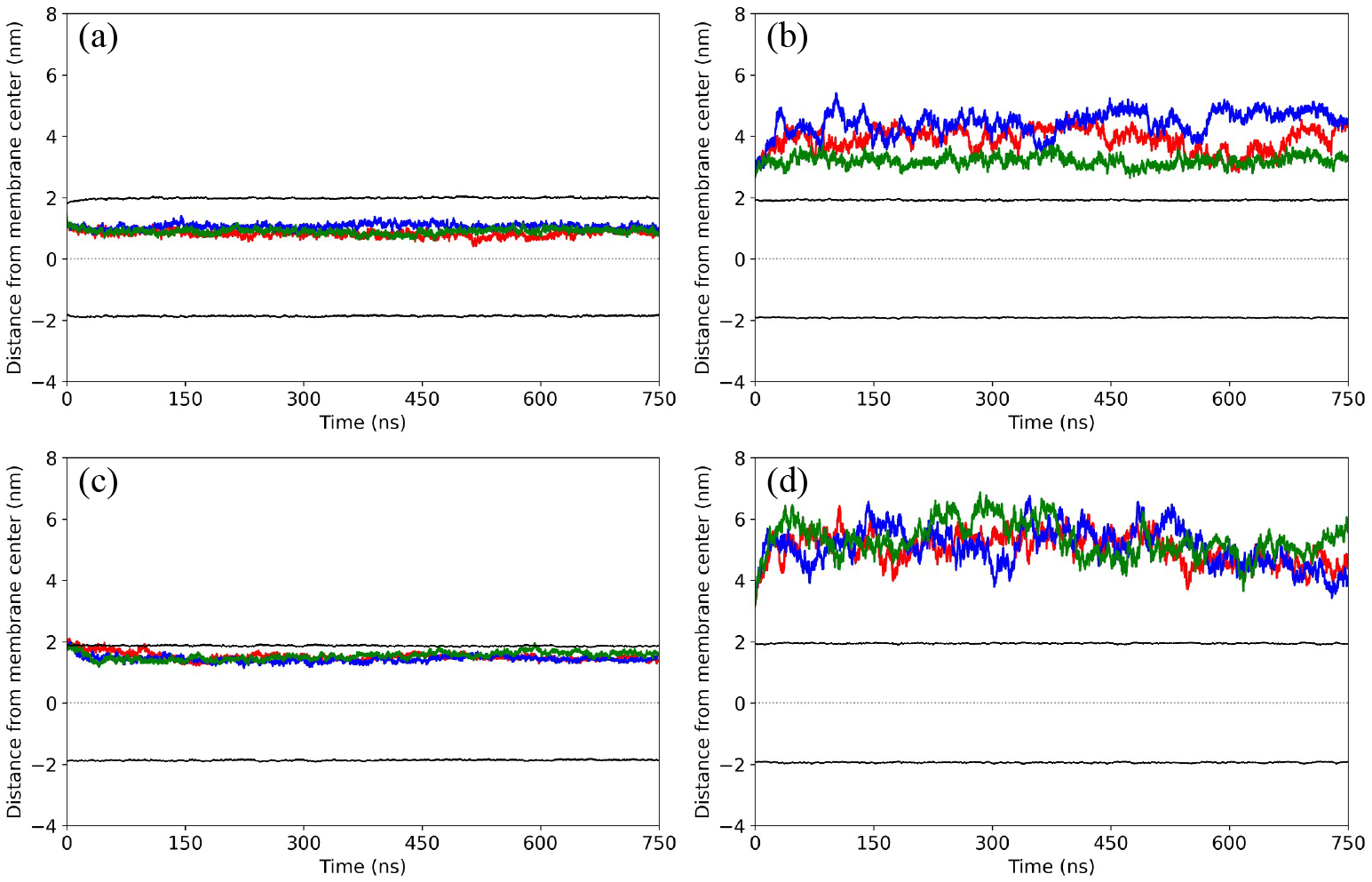
Positions along the z-axis of the COMs for melittin peptides and the phosphate groups of the two leaflets. Results are shown for systems: (a) system I-a, (b) system II-a, (c) system III-a, and (d) system IV-a. Black lines indicate the average z-position of phosphate groups in the upper and lower leaflets. Colored lines (red, blue, green) represent the COM of peptides from three independent simulations. In system I-a, the COM was located between the leaflets (approx. 1 nm from the bilayer center). In system III-a, peptides remain associated with the membrane surface. In contrast, for systems II-a and IV-a, peptides are in average 5 nm above the upper leaflet. No insertion was observed.

To understand why only simulations initiated from system I-a can successfully capture poration, we compared the initial configurations of the four systems. The membranes in systems II-a and IV-a remained intact, as shown in Figure S2 of the supplementary information. In contrast, both systems I-a and III-a had a hole (14 Å in diameter) on the upper leaflet. The hole was resulted from the partial embedding of melittins. Although the size of the hole in both cases are similar, only the one in system I-a spontaneously developed into a stable pore. Instead, the hole in system III-a rapidly disappeared during the simulations. To quantify this difference, we computed the number of water molecules located within *±*2.5 Å of the membrane center throughout the MD simulations. The three independent trajectories were concatenated for statistical analysis. For instance, probability distributions of number of water molecules were calculated using data from all three production runs. These probabilities were expressed as percentages of the total simulation time. This metric directly reflects the number of water molecules entering the pore during the simulations. Figure 5(a-b) presents the distribution of number of water molecules obtained from simulations of systems I-a and III-a. As expected, a large number of water molecules were observed in system I-a. This result is consistent with stable pore formation across three replicas. In contrast, very few water molecules were found at the bilayer center in system III-a, indicating the absence of pore.

**Fig. 5.**
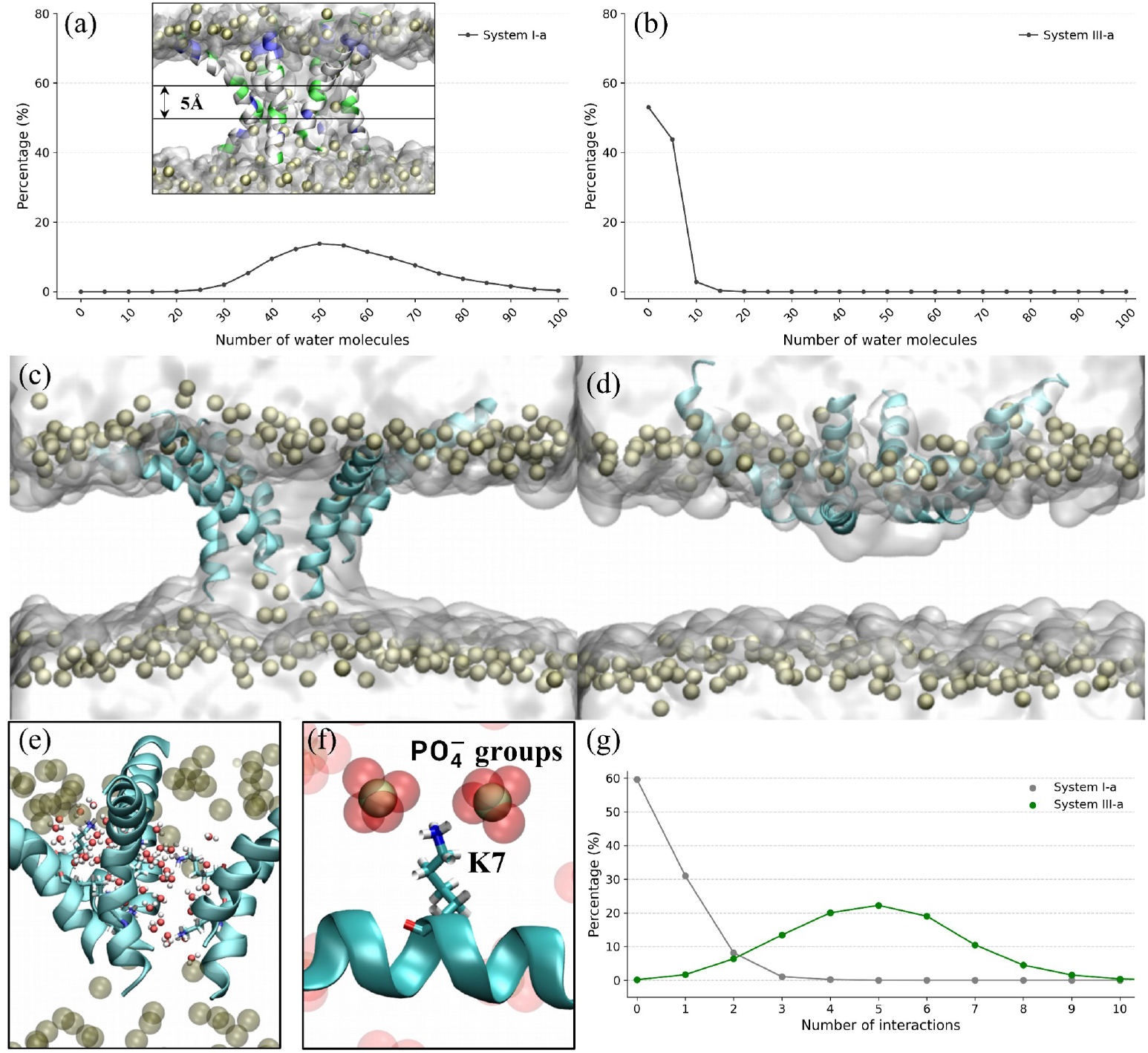
Water permeation and key peptide-lipid/water interactions. (a-b) The distribution of number of water molecules found at the membrane center (within *±*2.5 Å, z=0). The x-axis represents the number of water molecules and the y-axis shows the corresponding probability, expressed as a percentage of the total simulation length. Panels show results from: (a) system I-a, (b) system III-a. (c-d) Simulation snapshots at 2.5 ns, illustrating (c) transmembrane channel pore formation in system I-a and (d) the peptide S-state configuration in system III-a. (e-f) Zoomin of (c) and (d) to show K7’s interactions: (e) with nearby water molecules (within 3 Å) in system I-a, and (f) with phosphate groups in system III-a. (g) Distribution of number of phosphate groups in upper leaflet interacting with K7 in system I-a (gray) and system III-a (green). A phosphate group is considered to interact with K7 if it is within 5.5 Å of the K7 side chain’s 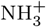 group.

Why did the initial hole on the upper leaflet of system I-a develop into a stable pore, while the hole in system III-a resealed during the simulations? A detailed comparison between Conf.I and Conf.III revealed that the orientation of cationic lysine-7 (K7) residue, differs significantly between the systems: in Conf.I, the K7 residues primarily face the solvent, whereas in Conf.III, they orient towards the negatively charged lipid phosphate groups. As shown in Figure 5, panels (c) and (d) display trajectory snapshots at 2.5 ns, which is the starting phase of the simulations, with system I-a forming a transmembrane pore and system III-a remaining in the S-state. Panels (e) and (f) further reveal distinct surroundings of K7: in system I-a, K7 primarily interacts with water molecules, while in system III-a, K7 strongly interacts with lipid phosphate groups. This observation is further supported by the number of phosphate groups interacting with K7. As shown in Figure 5(g), the number of phosphate groups interacting with K7 residues in system III-a ranges from 1 to 9, with a peak around 5, but K7 residues in system I-a are rarely in contact with membrane phosphate groups. It is clear that although an initial membrane defect can facilitate poration, strong electrostatic interactions between melittins and the lipid head groups can easily trap the peptides in an S-state, preventing pore formation.

### 3.2 Effects of Membrane Composition

Now, one may wonder, if the results of MD simulations (e.g., pore formation or not) depend so strongly on the initial configurations used, are the results still meaningful? To address this question, we extended the simulations to two different membrane models: the mammalian plasma membrane model (DOPC:Cholesterol, 9:1) and a Gram-negative bacterial membrane model (DOPE:DOPG, 3:1). These two models were selected for simulations because melittin-induced poration in the mammalian membrane model and the bacterial membrane model is associated with its hemolytic toxicity and antibacterial activity, respectively. The labels of the systems are provided in Table 1.

The distribution of number of water molecules, obtained from simulations of Conf.I with mammalian membrane model (system I-b) and of Conf.I with bacterial membrane model (system I-c), is depicted in Figure 6(a). In system I-b, the number of water molecules ranges from 5 to 80, with a peak occurring around 40. In contrast, the number of water molecules in system I-c peaks at 10. Both results indicate that pore formation process was observed in associated simulations, although the pore formed in the mammalian membrane model seems to be larger than that in the bacterial membrane model. Notably, the pore formed in the POPC bilayer is the largest among the three membrane models, as evidenced by its most probable water molecule count (50 molecules). Consistent with this observation, MD simulations demonstrate that the pore formed in the mammalian membrane model is comparatively smaller than that in the POPC bilayer. This finding is consistent with the widely recognized impact of cholesterol-induced membrane ordering [52, 53]. At the molecular level, cholesterols inserted among phospholipids can reduce the area per lipid and increase lipid order parameters. This ordering effect enhances bilayer cohesion, tightens lipid packing, and thereby raising the energetic barrier of transmembrane pore formation. Consequently, the presence of cholesterol significantly decreases membrane permeability [54, 55].

**Fig. 6.**
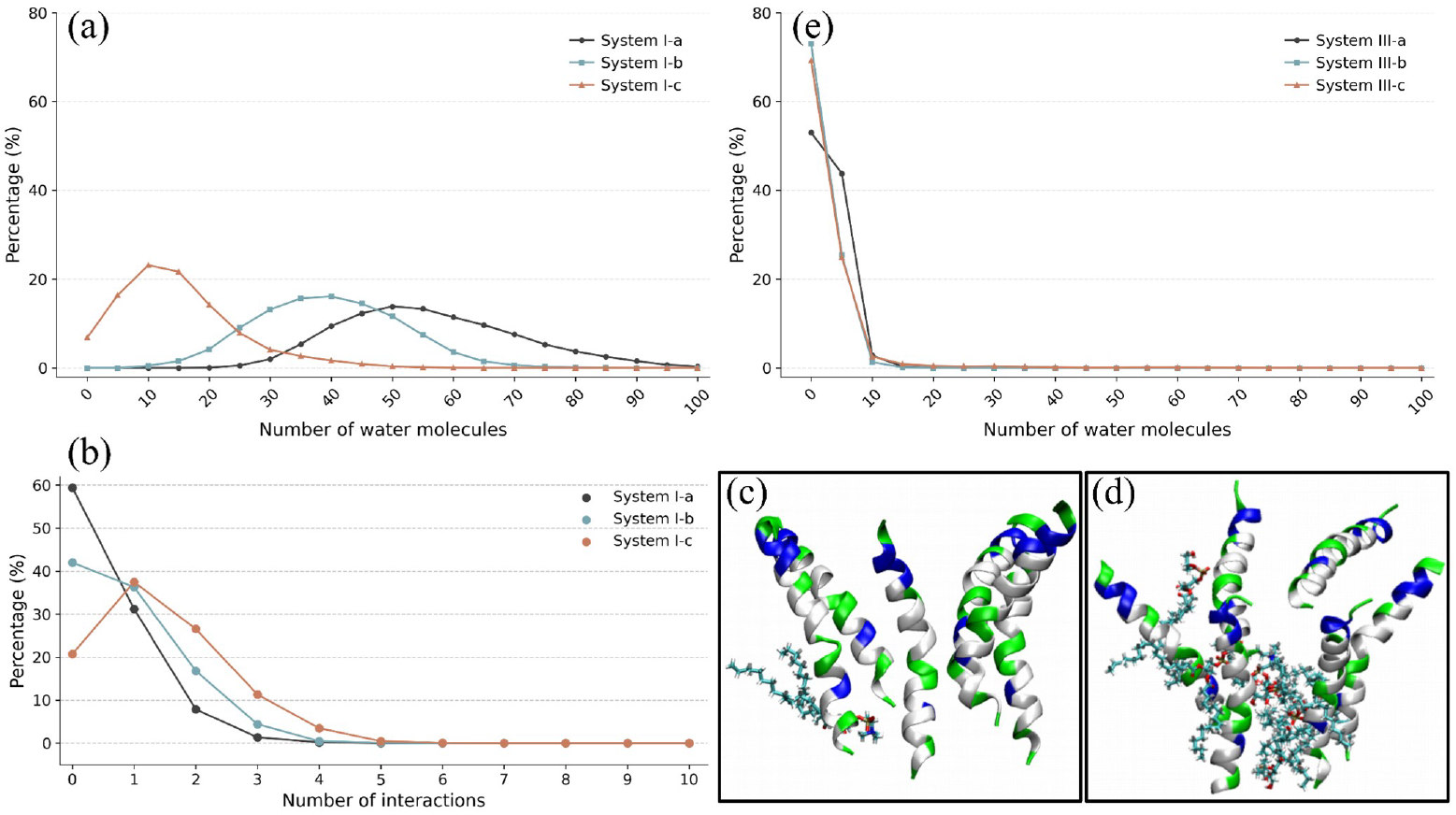
Analysis of water permeation and peptide-lipid interactions. (a) Distribution of number of water molecules found at the membrane center (within ±2.5 Å, z=0) for Conf.I. The x-axis represents the number of water molecules and the y-axis shows the corresponding probability, expressed as a percentage of the total simulation length. (b) Distribution of number of phosphate groups. A phosphate group is considered to interact with K7 if it is within 5.5 Å of the K7 side chain’s NH^+^ group. (c) Front view of peptide-lipid interactions in system I-b. (d) Front view of peptide-lipid interactions in system I-c. Peptides are shown in cartoon representation. Lipids with phosphates within 5.5 Å of K7s’ NH^+^ groups are displayed as bonds. (e) Distribution of number of water molecules found at the membrane center for Conf.III. Axes represent the same quantities as in panel (a).

Comparing the number of water molecules found in the bacterial membrane model and the mammalian membrane model leads to a rather unexpected conclusion: the pore formed in the bacterial membrane is much smaller than that in mammalian membrane. This observation may seem counterintuitive, especially when numerous studies highlight melittin’s affinity for bacterial cell membranes [56–58]. However, experimental data suggest otherwise. The minimum inhibitory concentration (MIC) of melittin against *E. coli* is 11.4 *μ*g/ml [59, 60], whereas the associated 50% hemolysis concentration (HC_50_) of human erythrocytes is 2.57 *μ*g/ml [61]. According to the data, melittin demonstrates a greater toxicity toward human erythrocytes compared 10 to its antibacterial activity against *E. coli*. Upon investigating the reason behind this simulation outcome, our attention was once again drawn to the K7 residues. In system I-c, these residues interact with phosphate groups more frequently than they do in systems I-a and I-b, as depicted in Figure 6(b). This interaction likely makes pore formation more challenging in system I-c. To be more precise, neutral DOPC lipids are rarely drawn to the peptides in system I-b (Figure 6(c)), but the strong electrostatic interactions between anionic DOPG and K7 can attract DOPG to block the pore, as shown in Figure 6(d). Lastly, Figure 6(e) indicates that no pore was formed in simulations initiated with Conf.III, irrespective of the membrane model used.

### 3.3 The Role of the Seventh Residue in Pore Formation

Having identified the impact of K7 on pore formation, we asked whether altering this residue could bias pore formation toward a particular membrane model. To test this hypothesis, two melittin mutants were simulated: one with K7 replaced by alanine (K7A) and the other by glutamine (K7Q). For both mutants, simulations were conducted using initial peptide configurations corresponding to Conf.I and Conf.III, across all three membrane models. The list of these systems was provided in Table 1. The distribution of water molecules for mutant simulations is depicted in Figure 7.

**Fig. 7.**
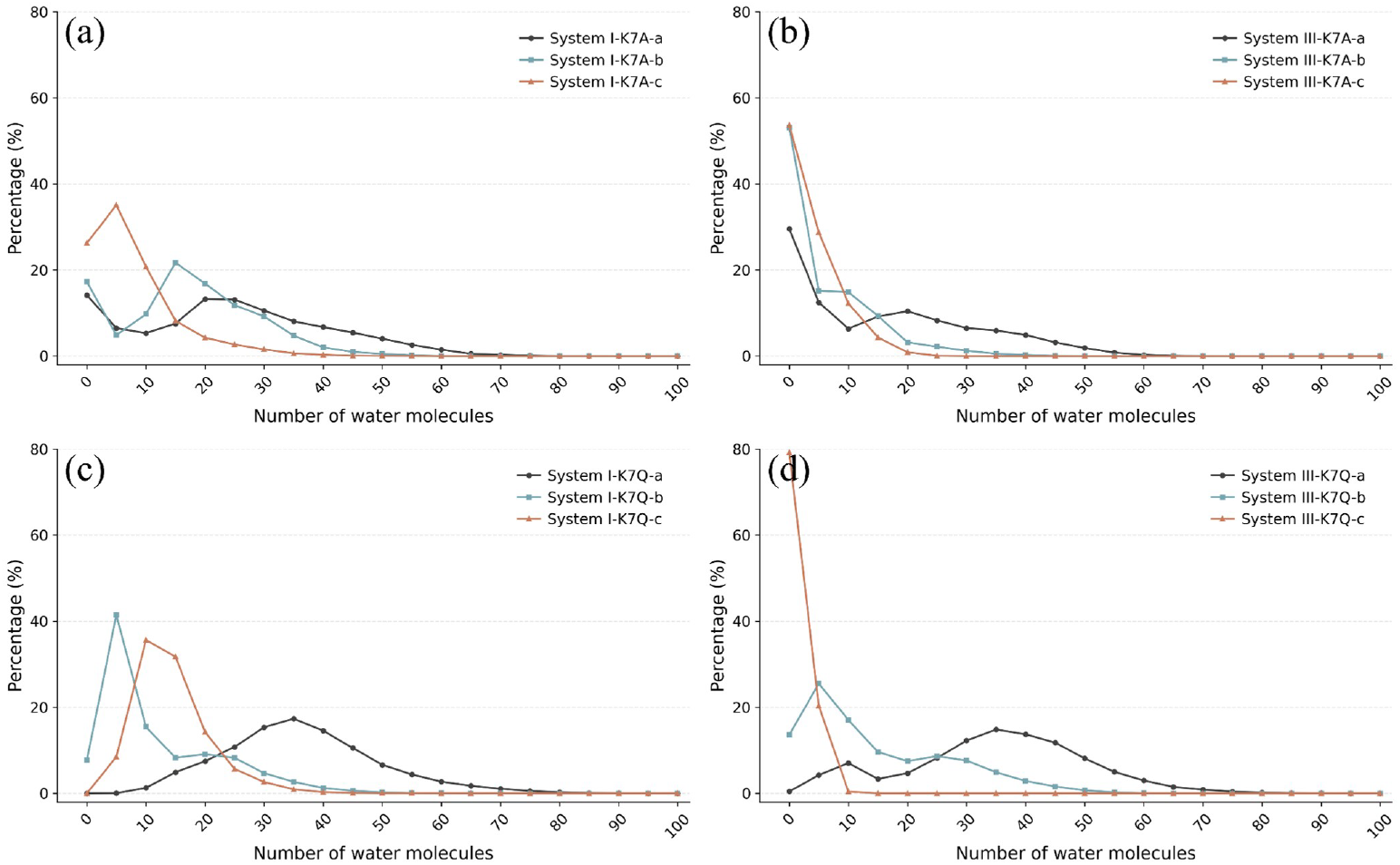
The distribution of number of water molecules found at the membrane center (within ±2.5 Å, z=0). The x-axis represents the number of water molecules and the y-axis shows the corresponding probability, expressed as a percentage of the total simulation length. Panels show results from: (a) Conf.I-K7A, (b) Conf.III-K7A, (c) Conf.I-K7Q, and (d) Conf.III-K7Q.

Panel (a) presents the results of K7A simulations with the initial peptide configuration arranged according to Conf.I. The three membrane systems are labeled as I-K7A-a, I-K7A-b, and I-K7A-c, for the POPC bilayer, the mammalian membrane model, and the bacterial membrane model, respectively. All three simulations demonstrate pore formation dynamics, as evidenced by the presence of water molecules in the membrane center. However, compared to the wild type melittin simulations, the pores formed by the K7A mutants appear less stable, and resulted in less water molecules to be found in the center of membrane.

Similarly, simulations of K7Q mutants with Conf.I’s initial configurations also captured the pore formation process, as indicated by the distribution of water molecules shown in panel (c). Notably, the pore formed in the mammalian membrane model (system I-K7Q-b) is now smaller than that formed in the bacterial membrane model (system I-K7Q-c). Apparently, replacing the cationic lysine with the polar glutamine reduced electrostatic interactions, while partially preserving the ability for water to pass. This allows the peptide to form pores more easily in bacterial membranes. In other words, mutating K7 to glutamine may reduce melittin’s toxicity towards human cells while maintaining its antimicrobial activity.

Interestingly, simulations conducted using Conf.III as the initial peptide configuration with K7A and K7Q mutants also exhibited pore formation. The distribution of number of water molecules of Conf.III-K7A and Conf.III-K7Q is shown in panels (b) and (d), respectively. After mutation, system III-K7Q-c showed no pore formation. While other systems exhibited transient pores, these structures were notably unstable and dissipated rapidly. This behavior differs from systems III-a, III-b, and III-c, where no pores formed at all. These observations suggest that Conf.III represents an unfavorable configuration for stable pore formation. This difference partly aligns with our understanding of K7 residues interaction with lipid phosphate groups, as neither alanine nor glutamine can anchor the peptides to the membrane surface through strong electrostatic interactions with the lipid headgroups. Notably, the pores formed in these simulations are transient and unstable. As demonstrated in Figure S3, the number of water molecules within 5 Å of the bilayer center gradually decays to zero as the simulation progresses. Nevertheless, these simulations demonstrate that while removing the positive charge at the seventh position facilitates peptide insertion and subsequent pore formation in some cases, the resulting configurations with Conf.III are largely unfavorable for stable pore formation.

## 4 Conclusions

In this study, we investigated the impact of initial configuration of melittins on poration dynamics using MD simulations. Four different initial peptide configurations and three distinct membrane models were examined. By counting the number of water molecules within ±2.5 Å of the membrane center, i.e., the number of water molecules in the pore, we assessed whether a stable transmembrane pore is formed during the simulations. Our results confirm that pore formation depends on the initial configuration, particularly on the presence of a small hole in the upper leaflet at the start of the simulation. However, not all small holes can develop into stable pores, as lysine at the seventh position on melittin (K7) can interact strongly with lipid phosphate groups, anchoring the peptides to the membrane surface. Consequently, simulations initiated with K7 facing lipids (Conf.III) only resulted in the S-state, even when a small hole was present initially. Thus, for melittin, only simulations starting from Conf.I successfully captured poration dynamics. Further analysis revealed that pore size decreases in the order: POPC bilayer *>* mammalian membrane model *>* bacterial membrane model. This trend aligns with melittin’s MIC against *E. coli* and its HC_50_ against human erythrocytes, and is consistent with the well-known effect of cholesterol-induced membrane ordering.

The role of the seventh residue on melittin was further examined by simulating two mutants. Substitution of K7 with alanine (K7A) abolished the strong electrostatic interaction between this residue and lipid phosphates, allowing pore formation even in simulations starting from Conf.III. However, the loss of lysine also reduced the peptide’s ability to attract water molecules from the solvent, resulting in smaller pores in simulations initiated from Conf.I. Notably, replacing K7 with glutamine (K7Q) reduced the interaction between the seventh residue and lipid phosphates while partially preserving its interaction with solvent water molecules. This led to an unexpected pore size distribution across the three membranes: POPC bilayer *>* bacterial membrane model *>* mammalian membrane model. While experimental validation is required to determine whether the MD simulations based on Conf.I can be used to guide the design of AMPs, such as to reduce toxicity or to increase antibacterial activity, our results demonstrate that all-atom MD simulations are sufficiently sensitive to reflect the difference in both membrane composition and residue mutations, suggesting that MD simulations is still a useful tool complementary to the widely use AI methods.

## 5 Data Availability

The data associated with this work are accessible through a Zenodo repository (https://doi.org/10.5281/zenodo.15241913). All molecular dynamics simulations were performed using the GROMACS software package (https://www.gromacs.org/). Analysis of simulation trajectories was conducted using the MDAnalysis Python library (https://www.mdanalysis.org/), and molecular visualization and figure preparation were done using VMD (Visual Molecular Dynamics) software (https://www.ks.uiuc.edu/Research/vmd/).

## 6 Author Contributions

Z.Z., Y.G., and Y.-C.C. contributed to conceptualization. Y.G., Z.Z., Y.L., and J.C.: investigation (MD simulations). J.C.: methodology, system building. P.X., J.G., L.Y., and C.-R.C.: validation. Y.G., Z.Z., and P.X.: formal analysis, writing original draft. T.-Y.L. and Y.-C.C.: supervision, project administration, funding acquisition.

## 8 Acknowledgments

The authors sincerely appreciate Kobilka Institute of Innovative Drug Discovery, The Chinese University of Hong Kong (Shenzhen) and National Yang Ming Chiao Tung University for financially supporting this research.

