## Supplementary Figures S1 to S3 for "Impact of Peptide Initial Configuration and Membrane Composition on Melittin’s Pore-Forming Ability under Unbiased All-Atom Molecular Dynamics Simulations"

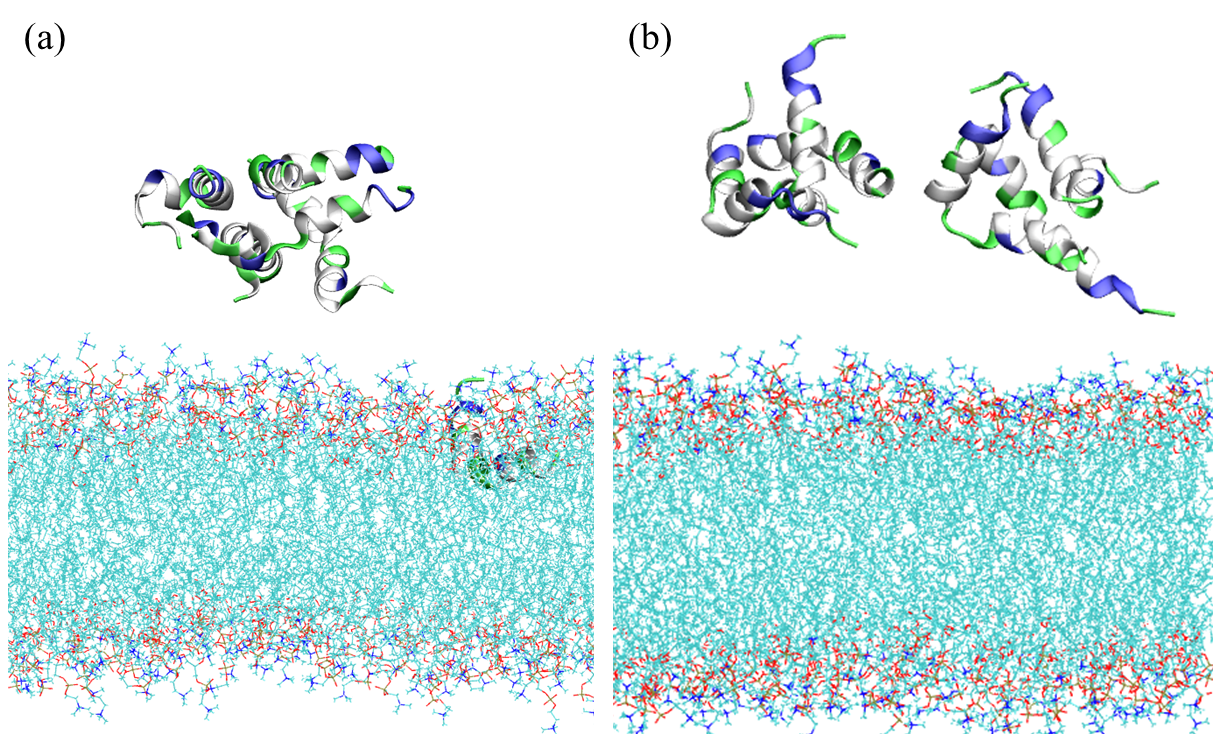

**Figure S1**: Representative snapshot of (a) System II-a (b) System IV-a, showing melittins remaining in the aqueous phase without pore formation.

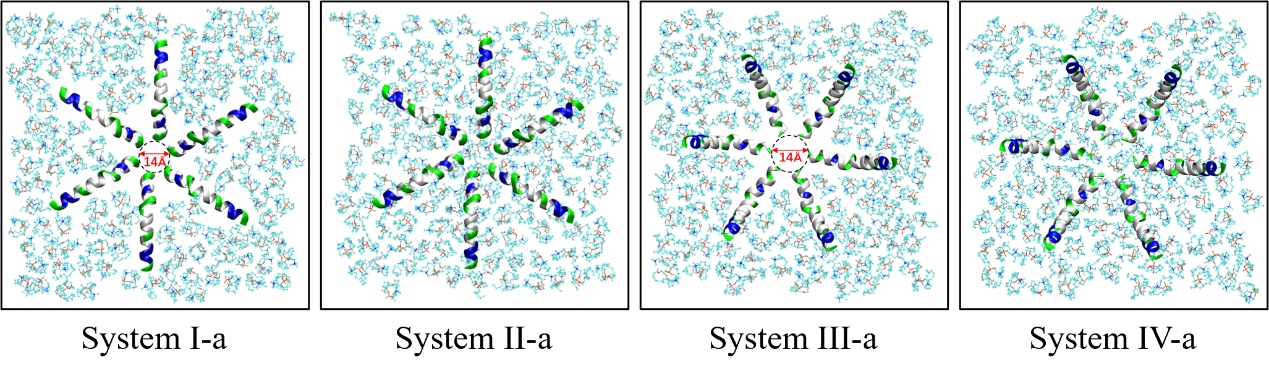

**Figure S2**: Top view of initial configurations showing the upper leaflet organization in different systems. System I-a and System III-a both exhibit a pre-formed hole of approximately 14 Å diameter due to the close peptide-membrane positioning. In contrast, System II-a and System IV-a show intact membrane surfaces with no initial hole. Peptides are shown in cartoon representation, with upper leaflet lipid molecules represented in line representation.

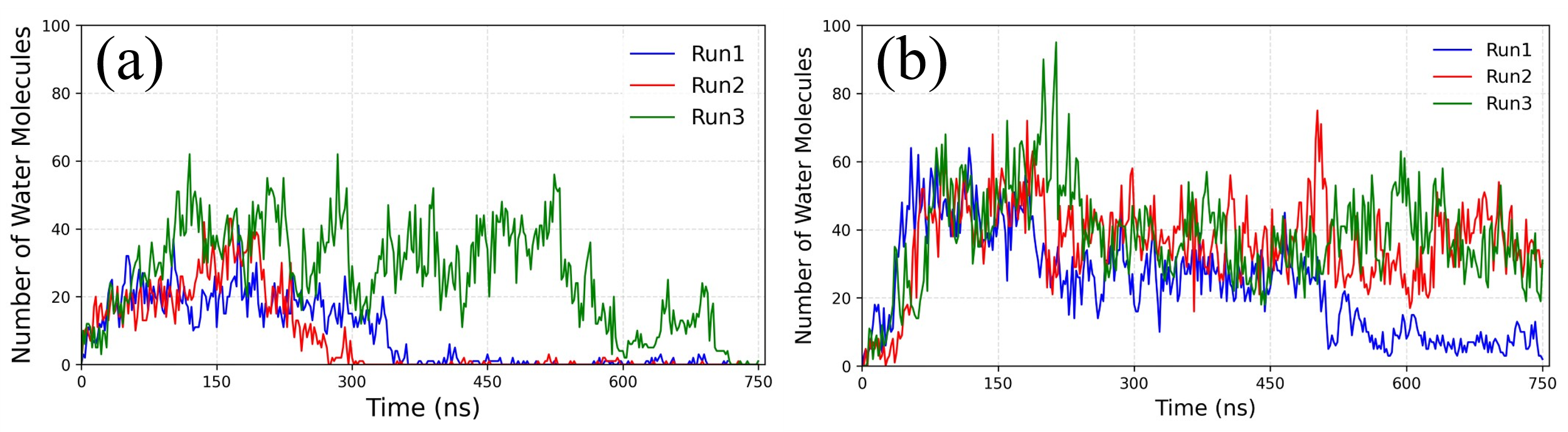

**Figure S3**: Number of water molecules within 5 Å of the bilayer center for three independent runs of (a) system III-K7A-a and (b) system III-K7Q-a. Data points sampled at 20-frame intervals are plotted against simulation time.
